# Mammarenavirus Z protein myristoylation and oligomerization are not required for its dose-dependent inhibitory effect on vRNP activity

**DOI:** 10.1101/2025.04.17.649453

**Authors:** Haydar Witwit, Juan Carlos de la Torre

## Abstract

We have recently documented the use of N-Myristoyltransferase inhibitors (NMTi) as an antiviral strategy against human pathogenic mammarenaviruses, including Lassa and Junin viruses, responsible for the hemorrhagic fever (HF) diseases Lassa fever (LF) and Argentine HF (AHF), respectively. Mammarenavirus Z matrix protein has been shown to exert a dose-dependent inhibitory effect in the activity of the virus ribonucleoprotein (vRNP) complex responsible for directing replication and transcription of the viral genome. Prevention of Z myristoylation by NMTi targeted Z protein for degradation, which resulted in inhibition of virus multiplication. Here, we review the recent findings of the use NMTi as antiviral, and reconcile the mechanistic process behind this inhibitory effect, more importantly, we used NMTi and G2A-mutated Z protein as a control of myristoylation function of wild type Z protein to elucidate that the vRNP suppression is monomeric-dependent activity of Z protein abundancy and it is independent of its myristoylation function.

## 1. Introduction

Several mammarenaviruses, chiefly Lassa virus (LASV) in Western Africa and Junin virus (JUNV) in the Argentine Pampas cause hemorrhagic fever (HF) diseases associated with high morbidity and mortality, posing important public health problems in their endemic regions. In addition, mounting evidence indicates that the worldwide-distributed mammarenavirus lymphocytic choriomeningitis virus (LCMV) is a neglected human pathogen of clinical significance in pediatric and transplantation medicine [1–3]. Current anti-mammarenavirus therapy is limited to an off-label use of ribavirin for which efficacy remains controversial [4]. Hence, the importance of developing novel therapeutics to combat human pathogenic mammarenaviruses.

Mammarenaviruses are enveloped viruses with a bi-segmented negative-stranded (NS) RNA genome [5]. Each genome segment, L (ca 7.3 kb) and S (ca 3.5 kb), uses an ambisense coding strategy to direct the synthesis of two polypeptides in opposite orientation, separated by a non-coding intergenic region (IGR). The L segment encodes the viral RNA dependent RNA polymerase (L), and Z matrix protein, whereas the S segment encodes the viral glycoprotein precursor (GPC) and the viral nucleoprotein (NP). GPC is co-and post-translationally processed to produce a stable signal peptide (SSP), and the mature GP1 and GP2 subunits that together with the SSP form the spikes that decorate the virus surface and mediate cell entry via receptor-mediated endocytosis [6–8]. GP1 mediates binding to the cellular receptor and GP2 the pH-dependent fusion event in the late endosome required for the release of the vRNP into the cell cytoplasm where it directs replication and transcription of the viral genome [9,10].

Early studies have shown that N-myristoylation is required for the role of Z protein in assembly and budding [11,12], and for the role of the SSP in the GP2-mediate fusion event [9]. These findings were based on the use of 2-hydroxy-myristic acid (2-HMA) and 2-HMA analogs as inhibitors of N-myristoyltransferase 1 (NMT1) and 2 (NMT2) responsible for catalyzing N-myristoylation in mammalian cells [13]. However, recent studies have demonstrated that 2-HMA acts off-target and does not inhibit N-myristoylation in a concentration range consistent with activity on NMT [14]. Therefore, we revisited the contribution of N-myristoylation in mammarenavirus infection using the validated on-target specific pan-NMTi (DDD85646 and IMP-1088) [15]. We found that DDD85646 and IMP-1088 exhibit very potent antiviral activity against LCMV and LASV in cultured cells. Cell-based assays probing different steps of the LCMV life cycle revealed that NMTi exerted its anti-LCMV activity by interfering with Z budding activity and GP2 mediated fusion [16] (Fig. 1). Our findings support the use of NMTi as a novel host-targeted antiviral strategy to combat LASV and other human pathogenic mammarenaviruses. Moreover, NMTi can be incorporated into combination therapy with direct acting antivirals. By targeting a host cell factor NMTi can pose a high genetic barrier for the selection of drug-resistant variants, a common problem with direct acting antiviral drugs. While Z dose-dependent suppression of vRNP activity is well documented [17–19], there is limited knowledge about the underlying mechanism [20–22]. Z protein has been shown to exhibit different degrees of oligomerization [16,23,24], whose biological implications remain poorly understood. Here we present evidence that Z oligomerization might be required for its budding activity, but not for its ability to inhibit vRNP activity.

**Figure 1.**
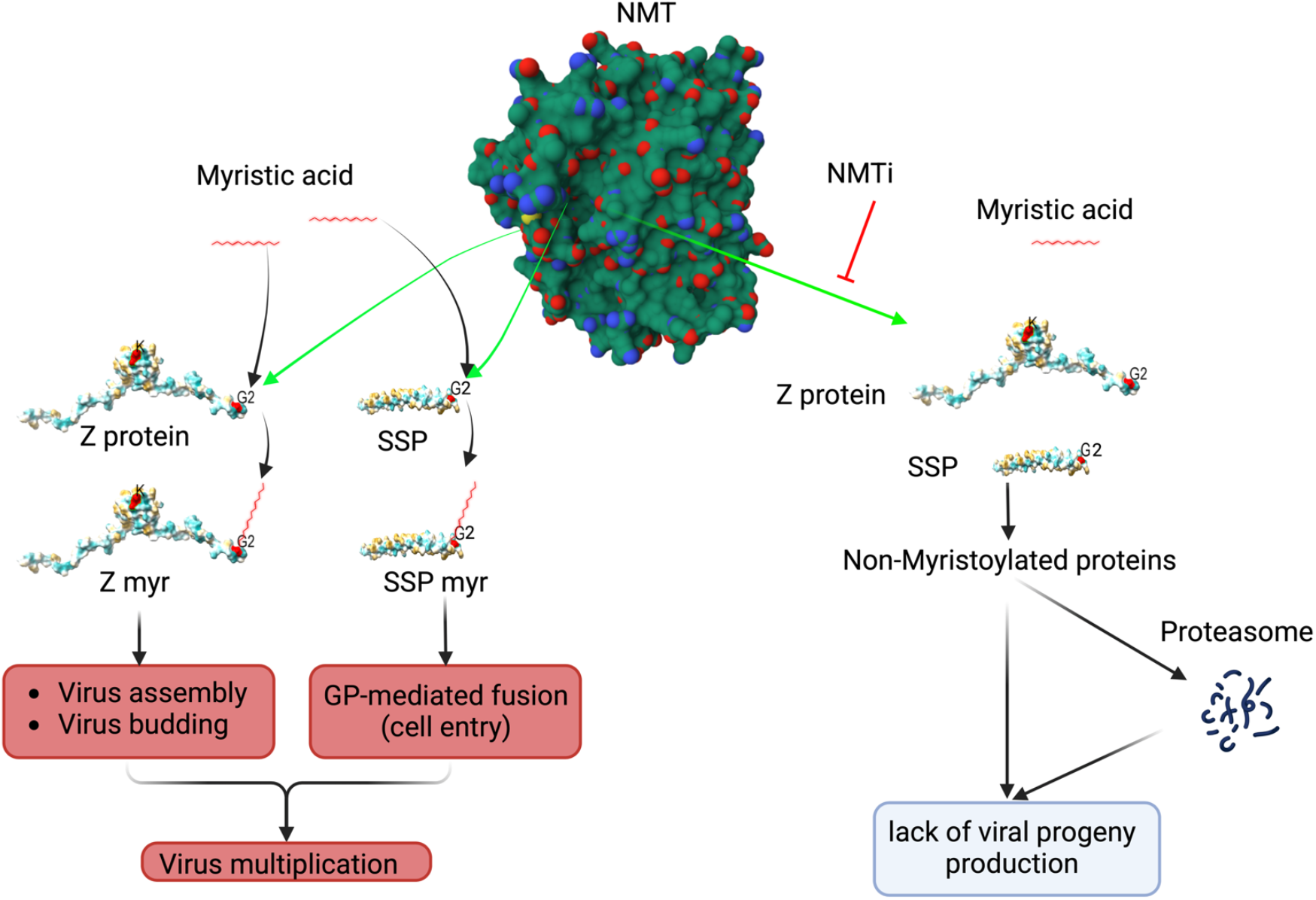
Proposed model of the effect of the NMT inhibitor on mammarenavirus cell entry and budding. NMT isozymes facilitate the addition of myristic acid to glycine (2) of SSP and Z protein, which protect them from proteasome mediated degradation. Myristoylated SSP interacts with GP2 to facilitate the fusion event in the late endosome required to complete the virus cell entry process, whereas myristoylated Z directs the virus assembly and budding process. Inhibition of SSP and Z myristoylation by NMT inh results in proteasome mediated degradation of SSP and Z which results in inhibition of virus multiplication. Z myr and SSP myr indicate myristoylated Z and SSP respectively. NMT pdb, 3IWE [25], and LASV Z matrix protein pdb, 2M1S [26], were used to generate the 3D figure using ChimeraX [27].

## 2. Materials and Methods

### 2.1 Cell line and Compounds

Human embryonic kidney cells stably expressing T antigen using SV40 promoter (HEK293T) were maintained in Dulbecco’s Modified Eagle’s Medium (DMEM) supplemented with 10% fetal bovine serum, 1mM penicillin and streptomycin, and 1 mM glutamine, the cells incubated at 37C and supplemented with 5% CO2. The NMTi IMP-1088, (5-[3,4-difluoro-2-[2-(1,3,5-trimethyl-1H-pyrazol-4-yl)ethoxy]phenyl]-N,N,1-trimethyl-1H-indazole-3-methanamine, Cat No. 25366-1) was purchased from Cayman Chemical, dissolved in methyl acetate at 11 mM, and kept in aliquots at −20. MG132 (MFG No. M7449) was purchased from Sigma-Aldrich (St. Louis, MO, USA).

### 2.2 Plasmids

The ORF of LCMV Z gene was cloned into pCAGGS and labeled with HA-tag, named here as Z-WT-HA, as described previously [16]. T7 polymerase in pCAGGS system and the S segment of LCMV flanked with T7 promoter at the amino terminus to express GFP. Both LCMV L polymerase and NP are expressed in pCAGGS systems.

### 2.3 Minigenome assay

HEK293 cells were seeded at 2×10^5^ per well in poly-L-lysine treated M12-well plates. Next day, cells were transfected with plasmids expressing LCMV L and NP proteins, the minimal trans-acting viral factors required for replication and transcription of the viral genome, and a plasmid directing intracellular synthesis of an LCMV minigenome directing expression of GFP (MG-GFP), together with a plasmid expressing the T7 RNA polymerase to launch primary synthesis of the MG-GFP, and incremental amounts of a plasmid expressing LCMV-Z tagged with HA (Z-WT-HA). At 18 h post-transfection, cells were treated with the indicated compounds, and at 24 h post-treatment, cells were fixed with 4% paraformaldehyde (PFA), and stained with anti-HA, followed by secondary fluorescence antibody to identify cells expressing the Z protein. The activity of the LCMV MG-GFP was assessed based on expression levels of GFP. Images were acquired at 4X magnification (BZ-X710 Keyence).

### 2.4 VLP assay

Viral like particle (VLP) production was done as described [11], briefly, HEK293T cells were transfected with pC_E as control, Z-WT or G2A mutant, and cell culture supernatant (CCS) collected at 48 or 72 hours post transfection (h pt). CCS were clarified by centrifugation (5 min, 5,000 RPM, 4°C). VLPs in clarified CCS were collected by ultracentrifugation (100,000 x *g*, 2 h, 4^0^C) through a 20% sucrose cushion in 50mM Tris-HCl pH7.5, 62.5 mM EDTA, 1% NP-40, 0.4% Na deoxycholoate. The pellet containing VLPs was resuspended in 60 µL of PBS and 20 µL of 4X Laemmli buffer added.

### 2.5 Immunoblotting

Protein concentration in samples was determined by BCA assay (Pierce™ BCA Protein Assay Kits, Cat# 23227, ThermoFisher, Waltham, MA USA). Samples (12 µg) were heated for 5 minutes at 95°C and separated by SDS-PAGE using a stain-free gel (Bio-Rad, Hercules, CA, USA). Total protein (TP) was detected after the activation of the stain free gel for one minute. Proteins were transferred to a low-fluorescence PVDF membrane (Bio-Rad, Hercules, CA, USA), and EveryBlot blocking buffer (Bio-Rad, Hercules, CA, USA) was used to block nonspecific antibody binding. Membrane was immunoreacted with an anti-HA antibody (Genscript, Piscataway, NJ, USA) overnight at 4^0^C. After three washes, the membrane was immunoreacted with a horseradish peroxidase (HRP) conjugated to anti-mouse antibody. Primary and secondary antibodies were diluted in OneBlock western-CL blocking buffer (Genesee Scientific, San Diego, CA). Protein bands were visualized with a chemiluminescent substrate (ThermoFisher Scientific).

### 2.6 Immunofluorescence Imaging

Images were acquired at 4X magnification using an BZ-X710 Keyence fluorescence microscope. Quantification and depiction of Z and GFP were done using BIOP JaCoP plugin, a colocalization quantification software, in FIJI imagej software (1.54f). The Otsu threshold setting was selected to quantify the degree of colocalization. The BIOP JaCoP generates, based on threshold setting, an automated figure that on the left side displays unmasked (top) and masked (bottom) panels that represent the Pearson correlation coefficient (X axis) and probability (Y axis) plots, and on the right side displays scatter plot representing the GFP (green, Y axis) and Z-HA (red, X axis) signals. The rationale to use BIOP Jacob plug is to quantify colocalization of GFP (a surrogate of the MG activity) and AlexaFLuor 568 red signal (a surrogate of Z matrix protein expression).

## 3. Results

### 3.1 Effect of NMTi on Z-mediated inhibition of vRNP activity

The Z protein has been shown to inhibit in a dose-dependent manner the activity of the mammarenavirus vRNP, responsible for directing replication and transcription of the viral genome, in cell-based minigenome (MG) assays. We have documented that treatment with NMTi target Z for degradation via the proteasome pathway [16]. We therefore predicted that Z inhibitory effect on the MG activity would be reduced in the presence of the pan-NMTi IMP-1088, and that treatment with the proteasome inhibitor MG132 would restore, in the presence of IMP-1088, the Z inhibitory effect on the MG activity. Using a cell-based LCMV minigenome (MG) system we found that treatment with the NMTi IMP-1088 counteract the Z-mediated inhibitory effect on the MG activity (Fig. 2). Treatment with the proteasome inhibitor MG132 restored expression levels of Z to those observed in the absence of IMP-1088, which resulted in the corresponding inhibitory effect on the activity of the LCMV MG (Fig. 2).

**Figure 2.**
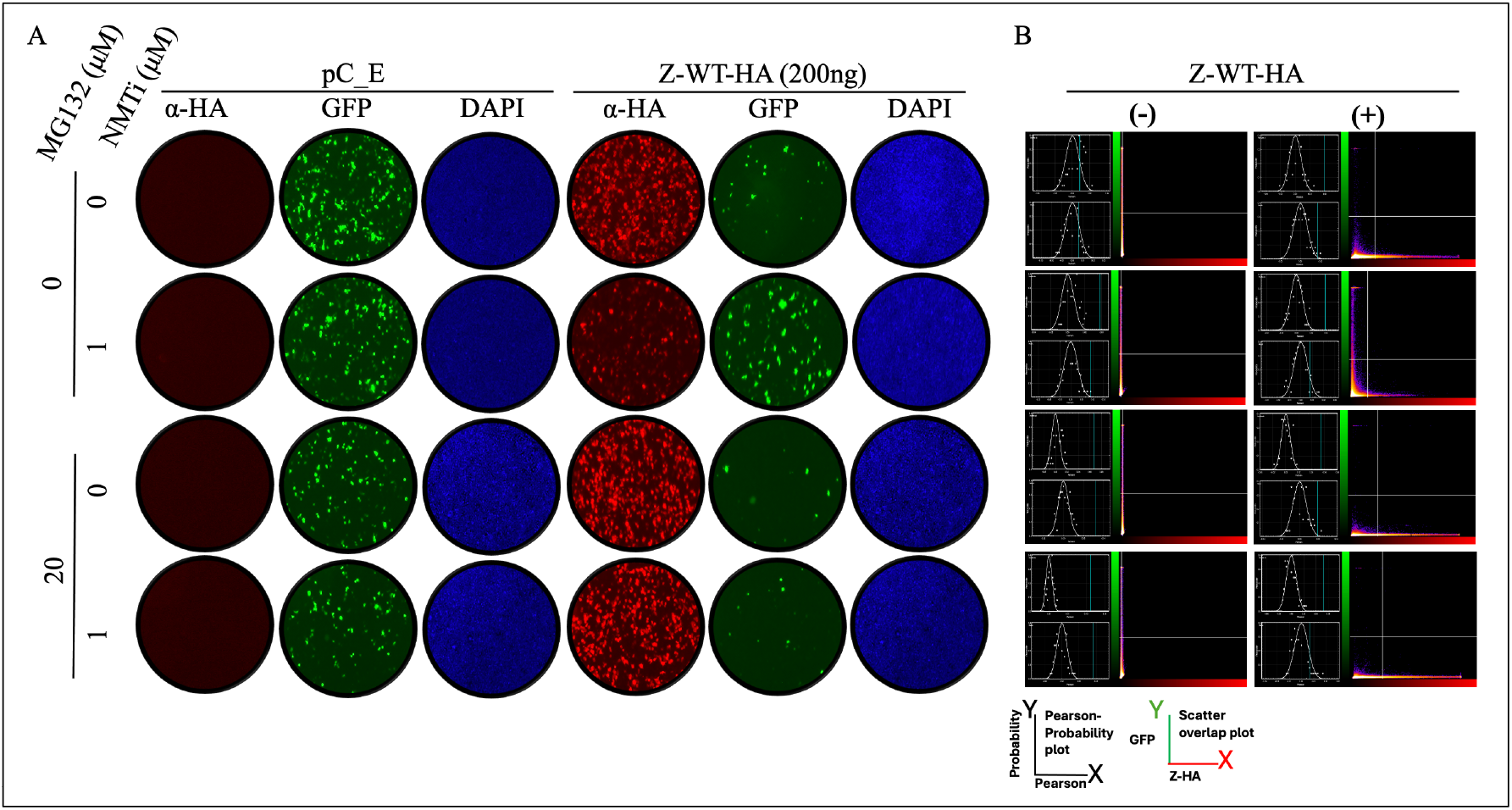
Effect of treatments with the NMTi (IMP-1088) and proteasome (MG132) inhibitors on the activity of the LCMV MG in the presence of LCMV WT Z protein. HEK293T cells were seeded at 2×10^5^ cells per well in poly-L-lysine treated M12-well plates. Next day, cells were transfected with pCAGGS plasmids expressing LCMV L and NP proteins, as well as T7RP, and a plasmid expressing the LCMV T7MG-GFP, together with 200 ng of a pCAGGS empty (pC-E) plasmid or expressing LCMV-Z-WT-HA. At 18 h post-transfection, cells were treated with the indicated compounds, and at 24 h post-treatment, cells were fixed with 4% PFA. Z-WT-HA expression was detected by IF using a rabitt polyclonal antibody to α-HA as primary antibody and an anti-mouse goat polyclonal antibody conjugated to Alexafluor 568 as secondary antibody. GFP was detected directly by epifluorescence. **A**. The activity of the LCMV MG-GFP was assessed based on expression levels of GFP. **B**. Left side of each panel displays unmasked (top) and masked (bottom) panels that represent the Pearson correlation coefficient (X axis) and probability (Y axis) plots, and on the right side displays scatter plot representing the GFP (green, Y axis) and Z-HA (red, X axis) signals. The BIOP Jacob plug quantified colocalization of GFP (a surrogate of the MG activity) and AlexaFLuor 568 red signal (a surrogate of Z matrix protein expression).

### 3.2 Role of N-myristoylation on Z-mediated inhibition of vRNP activity

Treatment with NMTi targeted Z protein for degradation, which prevented us from assessing whether myrisotylation was required for Z-mediated inhibition of vRNP activity. To address this question, we used the Z (G2A) mutant that cannot undergo N-terminal myristoylation. As predicted, treatment with IMP-1088 did not significantly affect expression levels Z (G2A) (Fig. 3). Notably, Z (G2A) inhibited MG-directed GFP expression with similar efficiency as Z WT (Fig. 3), indicating that myristoylation is not required for Z-mediated inhibition of the vRNP activity.

**Figure 3.**
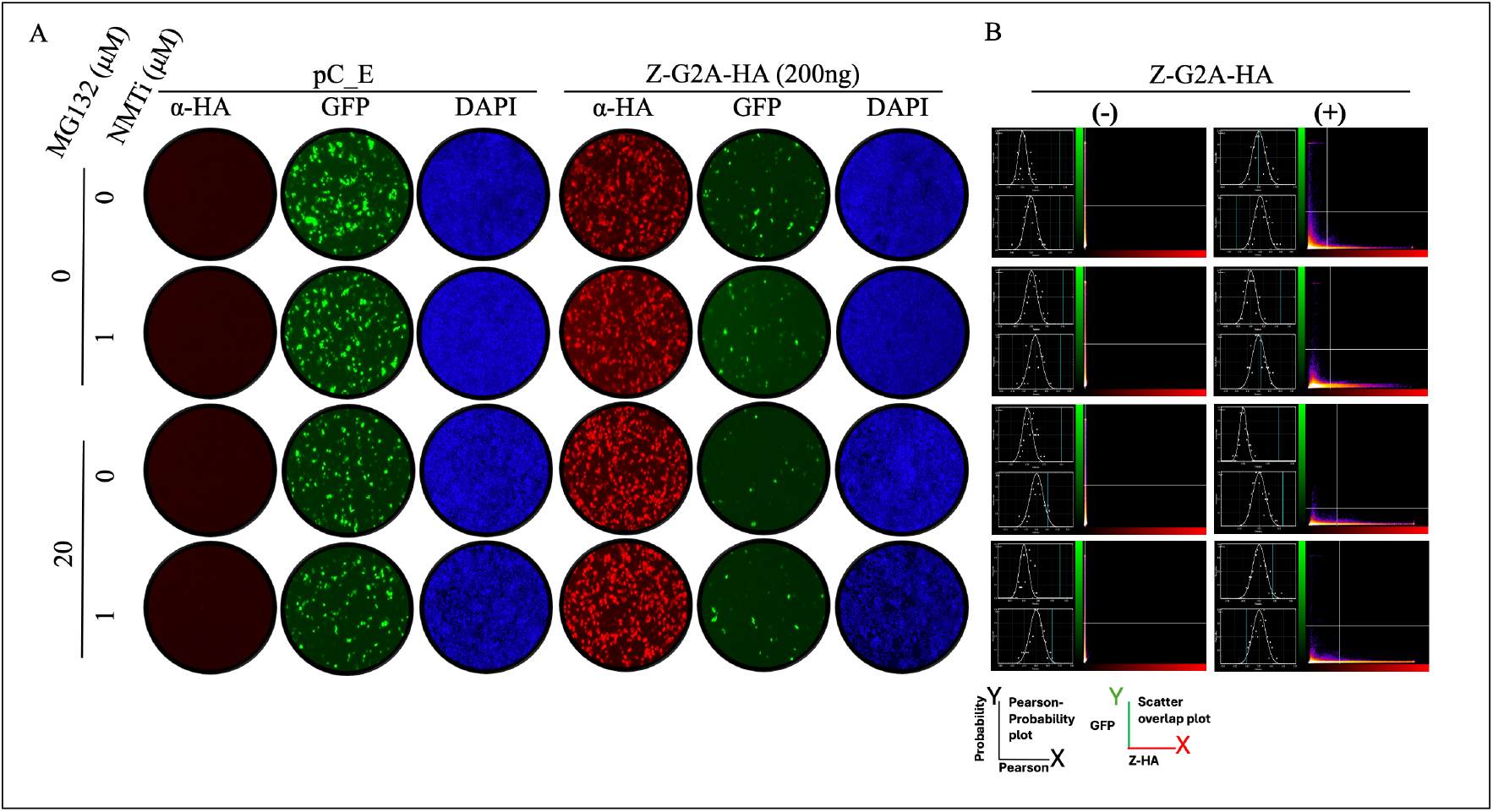
Effect of treatments with the NMT (IMP-1088) and proteasome (MG132) inhibitors on the activity of the LCMV MG in presence of LCMV Z G2A mutant. HEK293T cells were seeded at 2×10^5^ cells per well in poly-L-lysine treated M12-well plates. Next day, cells were transfected with pCAGGS plasmids expressing LCMV L and NP proteins, as well as T7RP, and a plasmid expressing the LCMV T7 MG-GFP, together with 200 ng of a pCAGGS empty (pC-E) plasmid or expressing LCMV-Z-G2A-HA. At 18 h post-transfection, cells were treated with the indicated compounds, and at 24 h post-treatment, cells were fixed with 4% PFA. Z-G2A-HA expression was detected by IF using a rabitt polyclonal antibody to α-HA as primary antibody and an anti-mouse goat polyclonal antibody conjugated to Alexafluor 568 as secondary antibody. GFP was detected directly by epifluorescence. **A**. The activity of the LCMV MG-GFP was assessed based on expression levels of GFP. **B**. Left side of each panel displays unmasked (top) and masked (bottom) panels that represent the Pearson correlation coefficient (X axis) and probability (Y axis) plots, and on the right side displays scatter plot representing the GFP (green, Y axis) and Z-G2A-HA (red, X axis) signals. The BIOP Jacob plug quantified colocalization of GFP (a surrogate of the MG activity) and AlexaFLuor 568 red signal (a surrogate of Z matrix protein expression).

### 3.3 Role of Z oligomerization on Z-mediated inhibition of vRNP activity and Z budding activity

We have documented the detection by western blotting of different oligomer species of Z protein in lysates of Z transfected cells [16]. We detected these Z oligomeric species in VLP produced in cells expressing WT, but not mutant G2A, Z protein (Fig. 4) mutant. In addition, rescuing the oligomer of G2A Z mutant, using proteasome inhibitor MG132, did not contribute to Z budding activity (Fig. 4B). These findings are consistent with published results showing the lack of Z oligomeric species in lysates and Z-containing VLP in cell culture supernatants of cells transfected with Z-G2A-HA [11,16]. We observed four Z species in VLP produced in cells transfected with Z-WT-HA with monomeric Z being the most abundant Z species. Inhibition of Z myristoylation using NMTi or mutation G2A results in degradation of Z oligomeric species [16], which complicates discerning the relative contributions of myristoylation and oligomerization to Z budding activity.

**Figure 4.**
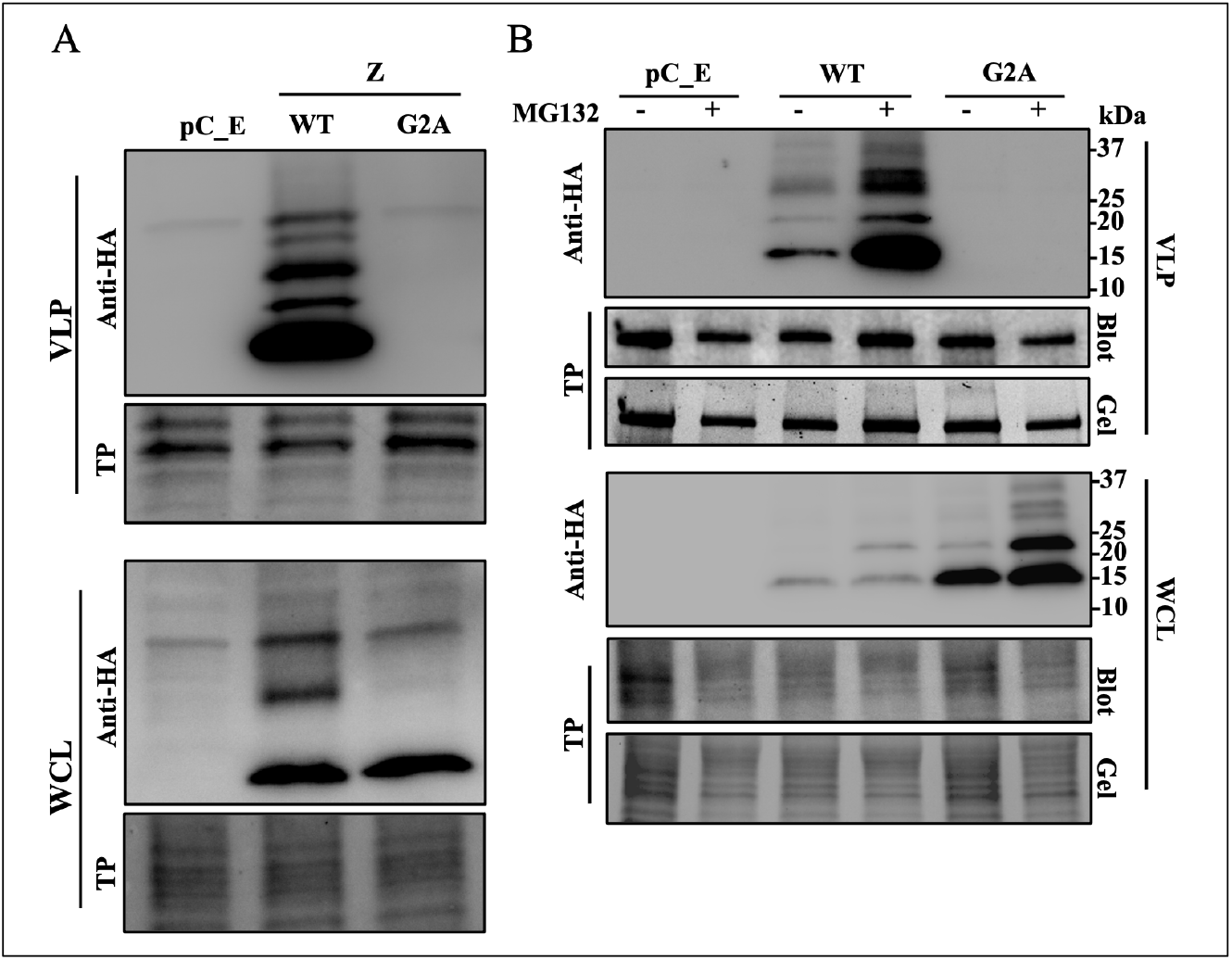
Oligomerization and cell egress of Z protein. **A**. HEK293T cells were seeded at 1×10^6^ cells per well in poly-L-lysine treated M6-well plates. Next day, cells were transfected with plasmids expressing LCMV Z WT or G2A mutant proteins tagged with HA tag or empty pCAGGS (pC-E). At 72 h post-transfection, supernatant and protein cell lysate were collected to extract VLP or whole cell lysate (WCL) protein respectively. **B**. HEK293T cells were seeded at 1×10^6^ cells per well in poly-L-lysine treated M6-well plates. Next day, cells were transfected with plasmids expressing LCMV Z WT or G2A mutant proteins tagged with HA tag or pC-E. Next day, media were aspirated and replaced with media containing MG132 (10 µg/mL) or vehicle control. At 48 h post-transfection supernatants were collected and whole cell lysate (WCL) prepared. VLP were collected by ultracentrifugation. Levels of Z in VLP and WCL samples were determined by western blotting using an antibody to HA. Membranes were imaged for total protein (TP) prior probing with anti-HA antibody.

## 4. Discussion

The mammarenavirus Z matrix protein plays critical roles in assembly and budding of matured infectious particles, processes that required Z myristoylation and oligomerization [11,24,28]. In addition, as with the matrix protein of other negative strand RNA viruses [29], Z has been shown to exhibit a dose-dependent inhibitory effect on mammarenavirus vRNP activity [18,19,30]. Biochemical and structural studies have indicated that Z can lock a polymerase-promoter complex [31], as well as induce conformational changes in L catalytic domains [20], which can account for Z-mediated inhibition of vRNP directed synthesis of viral RNA. These studies, however, did not address the question of whether myristoylation and oligomerization are required for Z-mediated inhibition of vRNP activity in infected cells.

Here we have documented that NMTi potent antiviral activity against LCMV and other mammarenaviruses correlated with proteasome mediated degradation of non-myristoylated Z protein, which disrupted virus particle assembly and budding thus resulting in reduced production of infectious progeny and restricted virus propagation [16] without inhibition of the vRNP activity. Treatment with the proteasome inhibitor MG132 in the presence of NMTi restored Z-mediated inhibition of vRNP activity (Fig 2), and the Z G2A mutant inhibited the activity of the LCMV MG with similar efficiency as Z WT (Fig 3). These findings indicate that Z myristoylation is not required for its inhibitory effect on vRNP activity. We also observed Z G2A exhibited highly reduced levels of oligomerization compared to Z WT (Fig 4), suggesting that oligomerization is not required for Z inhibitory effect on vRNP activity.

We have shown that dimers are the most abundant oligomer forms of Z in lysates from LCMV-infected or Z-transfected cells, and that these Z homodimers are efficiently targeted for degradation in the presence of NMTi, questioning the conclusion that Z homo-oligomerization is required for Z accumulation at the plasma membrane [24]. Our results also question that the G2 residue is required for oligomerization [24], since treatment with proteasome inhibitor MG132 resulted in similar expression levels of dimers of Z WT and Z G2A [16]. We propose that myristoylation at G2 prevents targeting of Z for degradation, thus allowing for the Z dose-dependent inhibitory effect on viral RNA synthesis mediated by the vRNP. Inhibition of Z myristoylation with NMTi results in proteasome mediated degradation of Z, thus preventing the Z inhibitory effect on vRNP activity, which can be restored by treatment with the proteasome inhibitor MG132 (Fig 5). We observed a prominent Z species of ~ 33 kDa, likely a trimer of Z, in cells transfected with Z WT that was absent in cells transfected with the G2A Z mutant and in the presence of MG132. The biological role of this Z species remains to be determined [16]. We also showed that oligomers of Z-G2A mutant lacked budding activity, further supporting that N-myristoylation is required for Z budding activity. It remains to be determined whether other protein lipidation modifications such as S-palmitoylation can substitute for N-myristoylation and facilitate Z association to membranes and budding.

**Figure 5.**
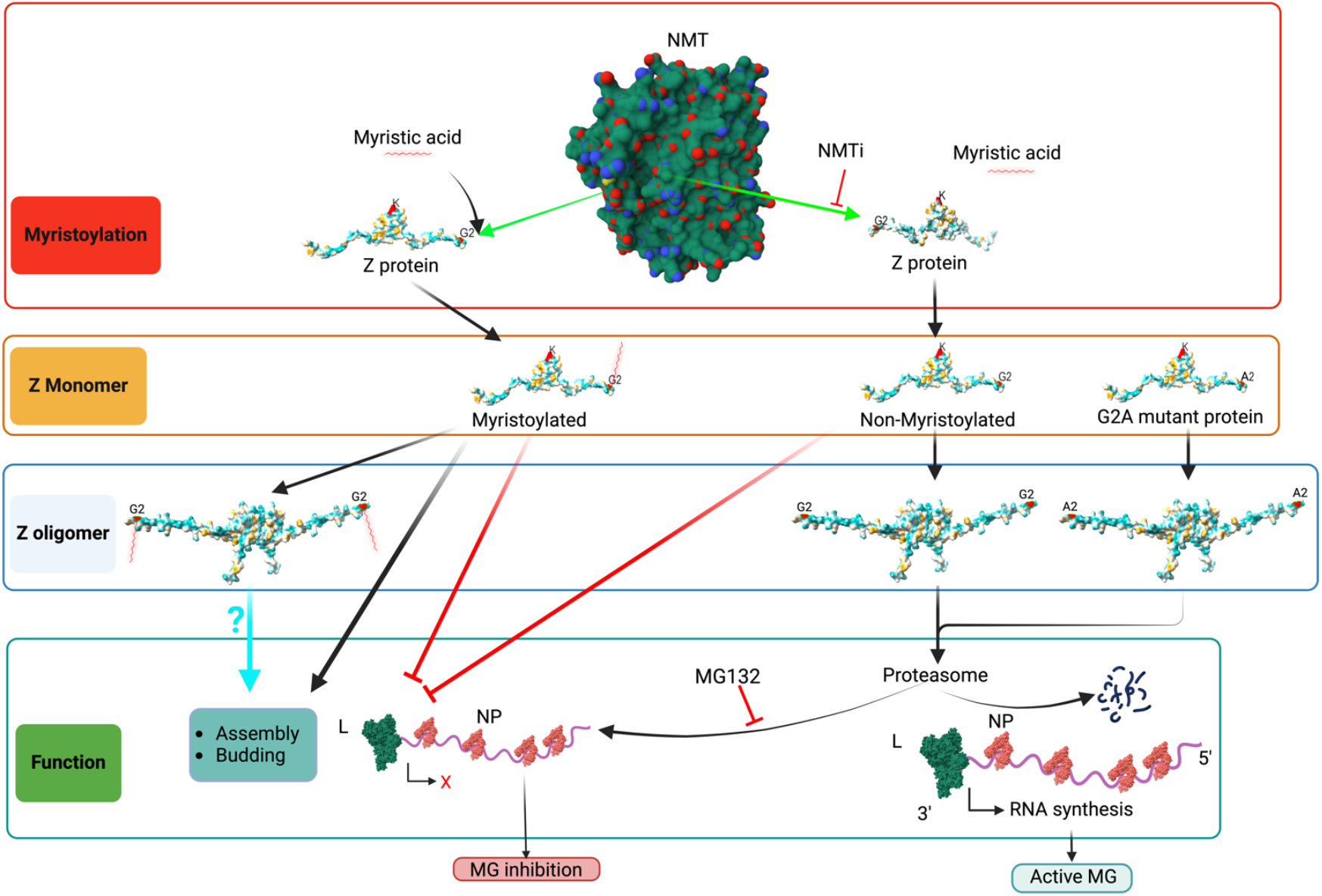
Proposed model of the effect of the NMT inh (IMP-1088) on the activity of the LCMV MG, budding, assembly and infectivity. NMT promotes Z myristoylation at G2, which prevents targeting of Z for degradation by the proteasome machinery, thus allowing for the Z dose-dependent inhibitory effect on viral RNA synthesis mediated by the vRNP and the consequent inhibition of MG directed GFP expression. Inhibition of Z myristoylation by IMP-1088 results in proteasome mediated degradation of Z, which will relief the Z inhibitory effect on the activity of the LCMV MG (right side of the graph). Treatment with the proteasome inhibitor MG132 prevents IMP-1088 induced Z protein degradation, thus resulting in LCMV MG inhibition. L polymerase (green), NP (red) and RNA (purple) are indicated. Red X indicates cessation activity, black angled arrow indicates RNA synthesis. Monomers of Z protein are sufficient to inhibit vRNP. Lack of Z myristoylation results in degradation of Z oligomers, which complicates the assessment of the role of Z oligomers in virus assembly, budding and infectivity. NMT pdb, 3IWE [25], and LASV Z matrix protein pdb, 2M1S [26], were used to generate the 3D figure using ChimeraX [27].

The use of NMTi to target host-cell lipidation processes opens the door to a new class of broad-spectrum anti-mammarenavirus therapy. Moreover, NMTi could be also incorporated into combination therapy strategies with direct acting antivirals, an approach expected to pose a high genetic barrier to the emergence of drug-resistant viruses, and to facilitate drug formulations with reduced toxicity [32]. Notably, the small molecule NMTi PCLX-001 has been shown to be safe and well tolerated in humans [33,34], supporting the interest of exploring the repurposing of NMTi to treat infections by human pathogenic mammarenaviruses. NMTi inhibitors have been shown to inhibit other viruses with myristoylated proteins, including picornaviruses [15] and vaccinia virus [32,35]. The use of specific pan-NMTi as a broad-spectrum antiviral strategy against viruses with myristoylated proteins warrants further investigation.

## Author Contributions

Conceptualization, HW and JCT; methodology, HW; software, HW; validation, HW, formal analysis, HW and JCT; investigation, HW.; resources, JCT; writing—HW; writing—review and editing, HW and JCT; visualization, HW; supervision, JCT; project administration, HW; funding acquisition, JCT. All authors have read and agreed to the published version of the manuscript.

## Institutional Review Board Statement

Not applicable

## Informed Consent Statement

Not applicable

## References

1. Yadav, K.; Mathur, G.; Ford, B.; Miller, R.; Group, C.W. A Case Cluster of Lymphocytic Choriomeningitis Virus Transmitted Via Organ Transplantation.: Abstract# D2381. Transplantation 2014, 98, 768.

2. Sayyad, L.E.; Smith, K.L.; Sadigh, K.S.; Cossaboom, C.M.; Choi, M.J.; Whitmer, S.; Cannon, D.; Krapiunaya, I.; Morales-Betoulle, M.; Annambhotla, P.; et al. Severe Non–Donor-Derived Lymphocytic Choriomeningitis Virus Infection in 2 Solid Organ Transplant Recipients. Open Forum Infectious Diseases 2025, 12, ofaf002, doi:10.1093/ofid/ofaf002.

3. Schafer, I.J.; Miller, R.; Ströher, U.; Knust, B.; Nichol, S.T.; Rollin, P.E.; Centers for Disease Control and Prevention (CDC) Notes from the Field: A Cluster of Lymphocytic Choriomeningitis Virus Infections Transmitted Through Organ Transplantation - Iowa, 2013. MMWR Morb Mortal Wkly Rep 2014, 63, 249.

4. Carrillo-Bustamante, P.; Nguyen, T.H.T.; Oestereich, L.; Günther, S.; Guedj, J.; Graw, F. Determining Ribavirin’s Mechanism of Action against Lassa Virus Infection. Sci Rep 2017, 7, 11693, doi:10.1038/s41598-017-10198-0.

5. Radoshitzky, S.R.; Buchmeier, M.; de la Torre, J.C. Emerging Viruses: Arenaviridae. In Fields Virology; Knipe, D. avid, Howley, P., Whelan, S., Eds.; 2020; Vol. I ISBN 978-1-975112-54-7.

6. Kunz, S.; Edelmann, K.H.; de la Torre, J.-C.; Gorney, R.; Oldstone, M.B.A. Mechanisms for Lymphocytic Choriomeningitis Virus Glycoprotein Cleavage, Transport, and Incorporation into Virions. Virology 2003, 314, 168–178, doi:10.1016/S0042-6822(03)00421-5.

7. Rojek, J.M.; Lee, A.M.; Nguyen, N.; Spiropoulou, C.F.; Kunz, S. Site 1 Protease Is Required for Proteolytic Processing of the Glycoproteins of the South American Hemorrhagic Fever Viruses Junin, Machupo, and Guanarito. Journal of Virology 2008, 82, 6045–6051, doi:10.1128/jvi.02392-07.

8. Fedeli, C.; Moreno, H.; Kunz, S. Novel Insights into Cell Entry of Emerging Human Pathogenic Arenaviruses. Journal of Molecular Biology 2018, 430, 1839–1852, doi:10.1016/j.jmb.2018.04.026.

9. York, J.; Nunberg, J.H. Role of the Stable Signal Peptide of Junín Arenavirus Envelope Glycoprotein in pH-Dependent Membrane Fusion. Journal of Virology 2006, 80, 7775–7780, doi:10.1128/jvi.00642-06.

10. York, J.; Nunberg, J.H. Myristoylation of the Arenavirus Envelope Glycoprotein Stable Signal Peptide Is Critical for Membrane Fusion but Dispensable for Virion Morphogenesis. Journal of Virology 2016, 90, 8341–8350, doi:10.1128/jvi.01124-16.

11. Perez, M.; Greenwald, D.L.; de La Torre, J.C. Myristoylation of the RING Finger Z Protein Is Essential for Arenavirus Budding. Journal of Virology 2004, 78, 11443–11448, doi:10.1128/jvi.78.20.11443-11448.2004.

12. Capul, A.A.; Perez, M.; Burke, E.; Kunz, S.; Buchmeier, M.J.; de la Torre, J.C. Arenavirus Z-Glycoprotein Association Requires Z Myristoylation but Not Functional RING or Late Domains. Journal of Virology 2007, 81, 9451–9460, doi:10.1128/jvi.00499-07.

13. Cordo, S.M.; Candurra, N.A.; Damonte, E.B. Myristic Acid Analogs Are Inhibitors of Junin Virus Replication. Microbes and Infection 1999, 1, 609–614, doi:10.1016/S1286-4579(99)80060-4.

14. Kallemeijn, W.W.; Lueg, G.A.; Faronato, M.; Hadavizadeh, K.; Grocin, A.G.; Song, O.-R.; Howell, M.; Calado, D.P.; Tate, E.W. Validation and Invalidation of Chemical Probes for the Human N-Myristoyltransferases. Cell Chemical Biology 2019, 26, 892–900.e4, doi:10.1016/j.chembiol.2019.03.006.

15. Ramljak, I.C.; Stanger, J.; Real-Hohn, A.; Dreier, D.; Wimmer, L.; Redlberger-Fritz, M.; Fischl, W.; Klingel, K.; Mihovilovic, M.D.; Blaas, D.; et al. Cellular N-Myristoyltransferases Play a Crucial Picornavirus Genus-Specific Role in Viral Assembly, Virion Maturation, and Infectivity. PLOS Pathogens 2018, 14, e1007203, doi:10.1371/journal.ppat.1007203.

16. Witwit, H.; Betancourt, C.A.; Cubitt, B.; Khafaji, R.; Kowalski, H.; Jackson, N.; Ye, C.; Martinez-Sobrido, L.; de la Torre, J.C. Cellular N-Myristoyl Transferases Are Required for Mammarenavirus Multiplication. Viruses 2024, 16, 1362, doi:10.3390/v16091362.

17. Cornu, T.I.; de la Torre, J.C. Characterization of the Arenavirus RING Finger Z Protein Regions Required for Z-Mediated Inhibition of Viral RNA Synthesis. J Virol 2002, 76, 6678–6688, doi:10.1128/jvi.76.13.6678-6688.2002.

18. Jácamo, R.; López, N.; Wilda, M.; Franze-Fernández, M.T. Tacaribe Virus Z Protein Interacts with the L Polymerase Protein To Inhibit Viral RNA Synthesis. J Virol 2003, 77, 10383–10393, doi:10.1128/JVI.77.19.10383-10393.2003.

19. López, N.; Jácamo, R.; Franze-Fernández, M.T. Transcription and RNA Replication of Tacaribe Virus Genome and Antigenome Analogs Require N and L Proteins: Z Protein Is an Inhibitor of These Processes. J Virol 2001, 75, 12241–12251, doi:10.1128/JVI.75.24.12241-12251.2001.

20. Liu, L.; Wang, P.; Liu, A.; Zhang, L.; Yan, L.; Guo, Y.; Xiao, G.; Rao, Z.; Lou, Z. Structure Basis for Allosteric Regulation of Lymphocytic Choriomeningitis Virus Polymerase Function by Z Matrix Protein. Protein Cell 2023, 14, 703–707, doi:10.1093/procel/pwad018.

21. Cornu, T.I.; de la Torre, J.C. RING Finger Z Protein of Lymphocytic Choriomeningitis Virus (LCMV) Inhibits Transcription and RNA Replication of an LCMV S-Segment Minigenome. J Virol 2001, 75, 9415–9426, doi:10.1128/JVI.75.19.9415-9426.2001.

22. Iwasaki, M.; de la Torre, J.C. A Highly Conserved Leucine in Mammarenavirus Matrix Z Protein Is Required for Z Interaction with the Virus L Polymerase and Z Stability in Cells Harboring an Active Viral Ribonucleoprotein. J Virol 2018, 92, e02256–17, doi:10.1128/JVI.02256-17.

23. Hastie, K.M.; Zandonatti, M.; Liu, T.; Li, S.; Woods, V.L.; Saphire, E.O. Crystal Structure of the Oligomeric Form of Lassa Virus Matrix Protein Z. J Virol 2016, 90, 4556–4562, doi:10.1128/JVI.02896-15.

24. Loureiro, M.E.; Wilda, M.; Levingston Macleod, J.M.; D’Antuono, A.; Foscaldi, S.; Buslje, C.M.; Lopez, N. Molecular Determinants of Arenavirus Z Protein Homo-Oligomerization and L Polymerase Binding?. J Virol 2011, 85, 12304–12314, doi:10.1128/JVI.05691-11.

25. Qiu, W.; Hutchinson, A.; Wernimont, A.; Lin, Y.-H.; Kania, A.; Ravichandran, M.; Kozieradzki, I.; Cossar, D.; Schapira, M.; Arrowsmith, C.H.; et al. Crystal Structure of Human Type-I N-Myristoyltransferase with Bound Myristoyl-CoA and Inhibitor DDD85646. To be published 2010, doi:10.2210/pdb3IWE/pdb.

26. Volpon, L.; Osborne, M.J.; Capul, A.A.; de la Torre, J.C.; Borden, K.L.B. Structural Characterization of the Z RING-eIF4E Complex Reveals a Distinct Mode of Control for eIF4E. Proc Natl Acad Sci U S A 2010, 107, 5441–5446, doi:10.1073/pnas.0909877107.

27. Meng, E.C.; Goddard, T.D.; Pettersen, E.F.; Couch, G.S.; Pearson, Z.J.; Morris, J.H.; Ferrin, T.E. UCSF ChimeraX: Tools for Structure Building and Analysis. Protein Science 2023, 32, e4792, doi:10.1002/pro.4792.

28. Perez, M.; Craven, R.C.; De La Torre, J.C. The Small RING Finger Protein Z Drives Arenavirus Budding: Implications for Antiviral Strategies. Proc. Natl. Acad. Sci. U.S.A. 2003, 100, 12978–12983, doi:10.1073/pnas.2133782100.

29. Baudin, F.; Petit, I.; Weissenhorn, W.; Ruigrok, R.W. In Vitro Dissection of the Membrane and RNP Binding Activities of Influenza Virus M1 Protein. Virology 2001, 281, 102–108, doi:10.1006/viro.2000.0804.

30. Cornu, T.I.; Feldmann, H.; de la Torre, J.C. Cells Expressing the RING Finger Z Protein Are Resistant to Arenavirus Infection. J Virol 2004, 78, 2979–2983, doi:10.1128/jvi.78.6.2979-2983.2004.

31. Kranzusch, P.J.; Whelan, S.P.J. Arenavirus Z Protein Controls Viral RNA Synthesis by Locking a Polymerase–Promoter Complex. Proceedings of the National Academy of Sciences 2011, 108, 19743–19748, doi:10.1073/pnas.1112742108.

32. Witwit, H.; Cubitt, B.; Khafaji, R.; Castro, E.M.; Goicoechea, M.; Lorenzo, M.M.; Blasco, R.; Martinez-Sobrido, L.; de la Torre, J.C. Repurposing Drugs for Synergistic Combination Therapies to Counteract Monkeypox Virus Tecovirimat Resistance. Viruses 2025, 17, 92, doi:10.3390/v17010092.

33. Sangha, R.; Davies, N.M.; Namdar, A.; Chu, M.; Spratlin, J.; Beauchamp, E.; Berthiaume, L.G.; Mackey, J.R. Novel, First-in-Human, Oral PCLX-001 Treatment in a Patient with Relapsed Diffuse Large B-Cell Lymphoma. Current Oncology 2022, 29, 1939–1946, doi:10.3390/curroncol29030158.

34. Sangha, R.S.; Jamal, R.; Spratlin, J.L.; Kuruvilla, J.; Sehn, L.H.; Weickert, M.; Berthiaume, L.G.; Mackey, J.R. A First-in-Human, Open-Label, Phase I Trial of Daily Oral PCLX-001, an NMT Inhibitor, in Patients with Relapsed/Refractory B-Cell Lymphomas and Advanced Solid Tumors. JCO 2023, 41, e15094–e15094, doi:10.1200/JCO.2023.41.16_suppl.e15094.

35. Priyamvada, L.; Kallemeijn, W.W.; Faronato, M.; Wilkins, K.; Goldsmith, C.S.; Cotter, C.A.; Ojeda, S.; Solari, R.; Moss, B.; Tate, E.W.; et al. Inhibition of Vaccinia Virus L1 N-Myristoylation by the Host N-Myristoyltransferase Inhibitor IMP-1088 Generates Non-Infectious Virions Defective in Cell Entry. PLoS Pathog 2022, 18, e1010662, doi:10.1371/journal.ppat.1010662.

